# Stemness prediction models reveal glioma aggressiveness and therapeutic targets in gliomas

**DOI:** 10.1101/2024.11.05.620942

**Authors:** Renan L.S. Simões, Maycon Marção, Emerson de Souza Santos, Sergio Akira Uyemura, Tathiane M. Malta

**Affiliations:** School of Pharmaceutical Sciences of Ribeirão Preto (FCFRP - USP), University of São Paulo, Ribeirão Preto, São Paulo, Brazil; Department of Developmental Neurobiology, St Jude Children’s Research Hospital, Memphis, TN USA

## Abstract

Gliomas are complex and heterogeneous primary brain tumors with a high degree of therapeutic resistance, particularly glioblastomas, which carry a poor prognosis. Cellular plasticity and stem cell-like features, or “stemness,” are increasingly recognized as key contributors to tumor progression and treatment resistance. In this study, we introduce two machine-learning-based prediction models designed to assess stemness in glioma samples using bulk gene expression data. One model captures fetal astrocyte characteristics (ASTsi), while the other identifies glioma stem cell traits (GSCsi). ASTsi was notably correlated with poor prognosis in IDHmut gliomas, whereas GSCsi was more indicative of stemness in IDHwt subtypes. Longitudinal data analysis showed that IDHwt and grade IV gliomas exhibit shifts in stemness indices upon recurrence, suggesting a phenotypic change linked to therapy resistance. Additionally, single-cell transcriptomic analysis confirmed that GSCsi can detect stem-like cell subsets in IDHwt gliomas. This approach enhances our understanding of glioma heterogeneity and reveals potential therapeutic targets.

## Introduction

Gliomas represent a significant portion of primary brain tumors and are known for their intricate heterogeneity and therapeutic resistance ^1^. Despite advances in understanding glioma biology, their etiology and oncogenic processes remain substantial challenges. Among the various types of gliomas, glioblastoma (GBM) is particularly notable due to its aggressiveness and poor prognosis, underscoring the urgent need for more effective therapeutic strategies ^2^.

Increasing evidence suggests that the plasticity of cells plays a significant role in the progression of tumors, their spread to other parts of the body, and their resistance to treatment; and characteristics resembling those of stem cells, commonly termed “stemness,” have been linked to unfavorable prognosis across various cancer types ^3,4^. The recognition of cancer stem cells (CSCs), including glioma stem cells (GSCs), has redefined our understanding of oncogenesis in brain tumors ^5^. GSCs are a distinct cellular subpopulation within gliomas, possibly responsible for tumor initiation, progression, heterogeneity, and treatment resistance ^6,7^. However, it is still unclear how GSCs originate and how they can be identified and characterized in tumor samples.

In this study, we propose an innovative approach to assess cancer stemness in glioma samples using machine learning algorithms trained on single-cell transcriptomic data. We present two distinct prediction models: one model based on non-tumor astrocytic signature and a second model based on glioma stem cell signature. When applied to bulk gene expression data from glioma patients, the models recapitulated aggressiveness and identified potential therapeutic targets. The prediction models applied to single-cell transcriptomic data allowed the identification of subsets of GSC.

## Results

### 1. Definition of inputs and creation of prediction models

#### 1.1. Fetal Astrocytes (ASTsi)

To build the prediction model, we accessed scRNA-seq data from 27 individuals (14 males and 13 females) of astrocytes derived from pluripotent stem cells from the work “Human Astrocyte Maturation Captured in 3D Cerebral Cortical Spheroids Derived from Pluripotent Stem Cells” (GSE99951) ^8^. After normalization and downstream analyses, we identified a cluster with high expression of fetal astrocyte marker genes (UBEC2C, NUSAP1, and TOP2A), as described in GSE99951 (FIGURE 1A, FIG SUPP 1A). We then used the expression data from this cluster to construct the fetal astrocyte prediction model. To create the model, we utilized the gelnet function from the gelnet package in R, which employs a One-Class Logistic Regression (OCLR) algorithm, as described in Malta 2018 ^4^. This machine learning approach allowed us to build a robust prediction model for fetal astrocytes based on their expression profile.

**Figure 1:**
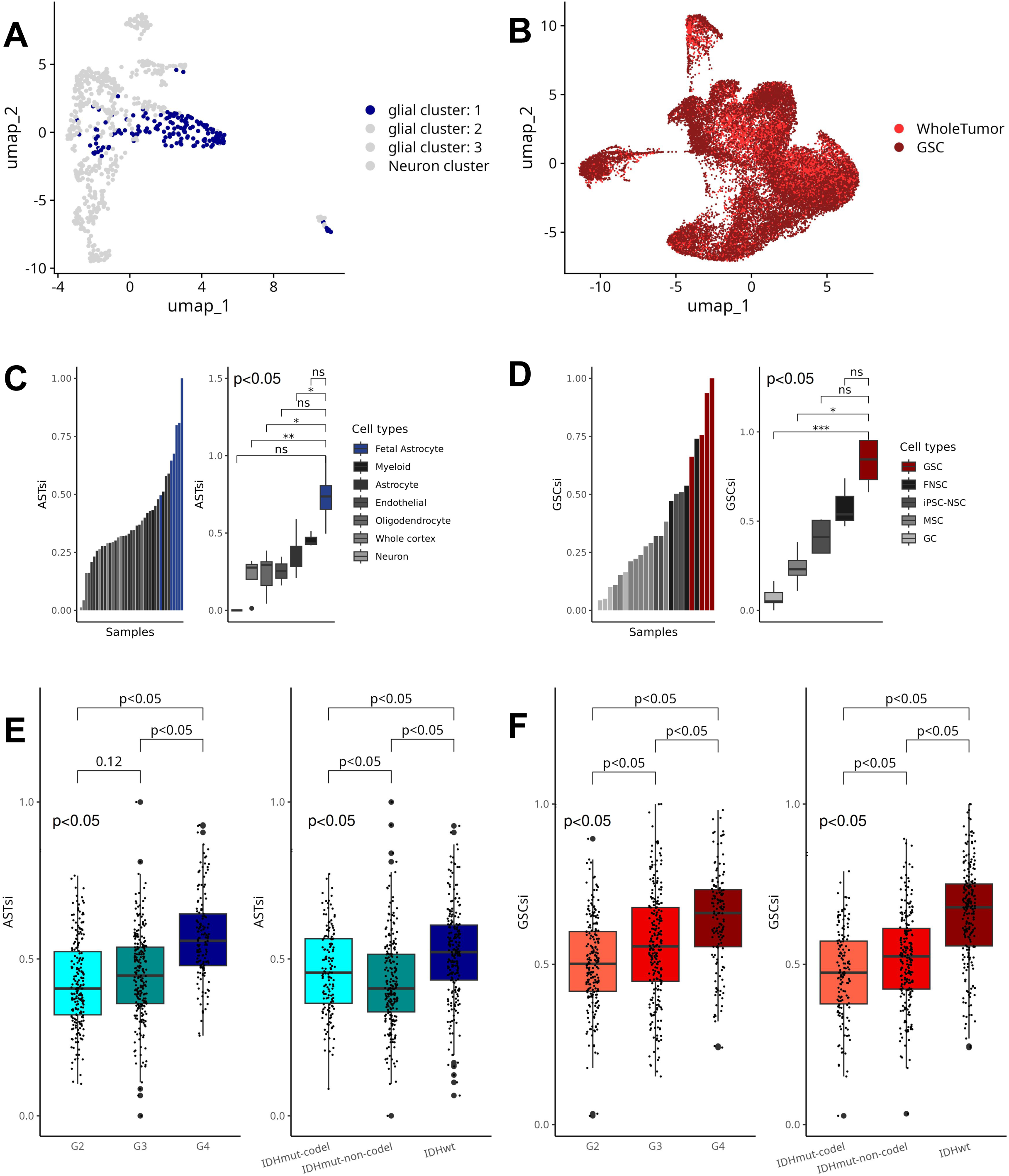
Characterization and Validation of Prediction Models and Stemness Indices. A) Uniform manifold approximation and projection (UMAP) plot of Human iPSC-derived scRNA-seq samples (n=707 scRNA-seq samples) highlighting in blue the samples defined as the fetal astrocyte cluster (n=171 scRNA-seq samples) B) UMAP plot of Human glioblastoma (n=7348 scRNA-seq samples) and derived GSCs (n=17868 scRNA-seq samples) scRNA-seq samples C) Barplot (left) and boxplot (right) showing the stemness indices (ASTsi) obtained in the validation dataset (n=41 bulk RNA-seq samples) D) Barplot (left) and boxplot (right) showing the stemness indices (GSCsi) obtained in the validation dataset (n=24 bulk RNA-seq samples) E) Boxplots showing the distribution of ASTsi in LGG/GBM data from TCGA (n=600 bulk RNA-seq samples). On the left showing the distribution of ASTsi separating by tumor grade (n=214 G2, 241 G3, 145 G4), on the right showing the distribution of ASTsi separating by IDH mutation status (n=151 IDHmut-codel, 226 IDHmut-non-codel, 223 IDHwt). F) Boxplots showing the distribution of GSCsi in LGG/GBM data from TCGA. On the left showing the distribution of GSCsi separating by tumor grade, on the right showing the distribution of GSCsi separating by IDH mutation status.

#### 1.2. Glioma Stem Cells (GSCsi)

The GSC model construction was performed using scRNA-seq gene expression data from derived GSC samples of 3 patients containing both whole tumor and GSC from the study “Single-cell RNA-seq reveals that glioblastoma recapitulates a normal neurodevelopmental hierarchy” ^9^. Data from the 3 patients were integrated to enable comparative and integrative analyses. After clustering, it was observed in the UMAP that GSCs exhibit a gene expression profile similar to whole tumor samples of gliomas (FIGURE 1B, FIG SUPP 1B). We then selected the expression data of GSCs from the 3 patients as input for the GSC prediction model. Similar to the ASTsi model, we employed the gelnet function from the gelnet package in R, applying the OCLR algorithm to construct the GSC prediction model. This model is designed to capture stem cell-like properties based on gene expression data, reflecting the unique characteristics of glioma stem cells.

#### 1.3. Validation of the models

We then performed validation of the prediction models with external data. Gene expression data containing different cell types, including samples similar to those used as input for the models, namely fetal astrocytes and GSCs, were accessed. Both prediction models proved effective in detecting populations with higher indices - ASTsi for the fetal astrocyte model and GSCsi for the GSC model. The fetal astrocyte prediction model was able to identify fetal astrocytes with higher indices and to separate fetal astrocytes from mature astrocytes (FIGURE 1C). The GSC model also generated higher indices for the GSCs in the dataset and was able to separate GSCs from glioma cells (FIGURE 1D).

#### 1.4. Histopathological and molecular characteristics associated with Stemness

We applied the prediction models generated for fetal astrocytes (ASTsi) and glioma stem cells (GSCsi) to gene expression samples of gliomas (LGG and GBM) from The Cancer Genome Atlas (TCGA), generating stemness indices for each tumor sample. These indices provided a quantitative measure of the stem cell-like properties within the tumors. We considered the main histopathological and molecular characteristics of gliomas, namely tumor grade and IDH mutation status, to biologically understand the distribution of stemness indices generated by the prediction models. The most aggressive subtypes (IDHwt and Grade 4) exhibited a higher presence/enrichment of tumor stem cells according to the obtained indices (FIGURE 1E, FIGURE 1F). These results support our hypothesis that tumors with more aggressive characteristics present a higher presence of cells with a tumor stem cell phenotype. To confirm that the generated indices reflect the presence of tumor stem cells, we applied the pluripotent stem cell prediction model (mRNAsi.2018), built by Malta in 2018 ^4^, to the same data. The model constructed with pluripotent stem cells is unable to stratify the tumor samples considering grade and IDH status (FIG SUPP 1C) in a biologically expected manner. In the distribution by IDH status, it was observed that IDHmut-codel had higher mRNAsi.2018 than IDHwt and IDHmut-noncodel, which does not reflect biological behavior of these tumors. These results indicate that these tumor stem cells likely have a certain degree of stemness but are not as undifferentiated as pluripotent stem cells. It is more likely that GSCs have a profile of intricate stem cells with tissue-specific characteristics, which can be confirmed by the correlation analysis between the obtained indices (ASTsi, GSCsi, and mRNA2018) with the models (FIG SUPP 1D).

### 2. ASTsi and GSCsi associated with prognosis

Considering the ability of the prediction models to identify glioma samples from TCGA by grade and IDH status, we proceeded with overall survival (OS) and progression-free survival (PFS) analyses to verify if our prediction models were associated with patient survival data (FIGURE 2). The TCGA cohort was separated into IDHmut and IDHwt samples to avoid survival bias, given that our models were capable of separating samples by IDH mutation status, and there is a significant difference in prognosis between the two groups. The results provide evidence that greater stemness, specifically that generated by the fetal astrocyte model (ASTsi) is related to the worse prognosis of patients with IDHmut gliomas. For both OS and PFS, IDHmut samples showed a greater risk association (Hazard Ratio) with ASTsi. Although the CI 95% and p.value found for GSCsi in the IDHmut samples do not present evidence of a risk association, the OSHR found was significantly higher than that found in the IDHwt samples. In the IDHwt samples, no evidence of risk association with the indices obtained by the models was found (FIGURE 2A). These results suggest that although IDHwt gliomas have higher stemness indices than IDHmut (as seen in Figure 1), the risk association with stemness is more evident in IDHmut gliomas, especially stemness characterized by the fetal astrocyte phenotype. We then divided the TCGA samples into two qualitative groups of high and low stemness according to ASTsi or GSCsi. The top 25% of samples with the highest ASTsi or GSCsi (above the 3rd quartile) were defined as high stemness, while the bottom 25% of samples with the lowest ASTsi or GSCsi (below the 1st quartile) were defined as low stemness. Overall, the qualitative division of stemness did not provide evidence of a difference in patient survival related to GSCsi. On the other hand, the division into low and high ASTsi showed evidence of two groups with different prognoses in IDHmut glioma samples (FIGURE 2B). Among IDHmut gliomas, there are other histological and molecular subclassifications, including stratification based on methylation clusters defined by Cecarelli in 2016, which proved efficient in identifying different prognoses ^10^. The categorical division into high and low stemness observed in the fetal astrocyte model proved efficient in identifying distinct prognoses. Therefore, we used ASTsi to separate IDHmut samples into high and low stemness based on DNA methylation groups (G-CIMP-low, G-CIMP-high, and Codel) (FIGURE 2C). The G-CIMP-low group with high ASTsi showed the worst prognosis, as anticipated, being the most aggressive subtype with the poorest survival. The separation into high and low stemness in the G-CIMP-high and Codel groups showed two distinct groups with different prognoses among IDHmut glioma samples, indicating that the ASTsi stratify glioma samples with the worst prognosis even within the same DNA methylation subtype.

**Figure 2:**
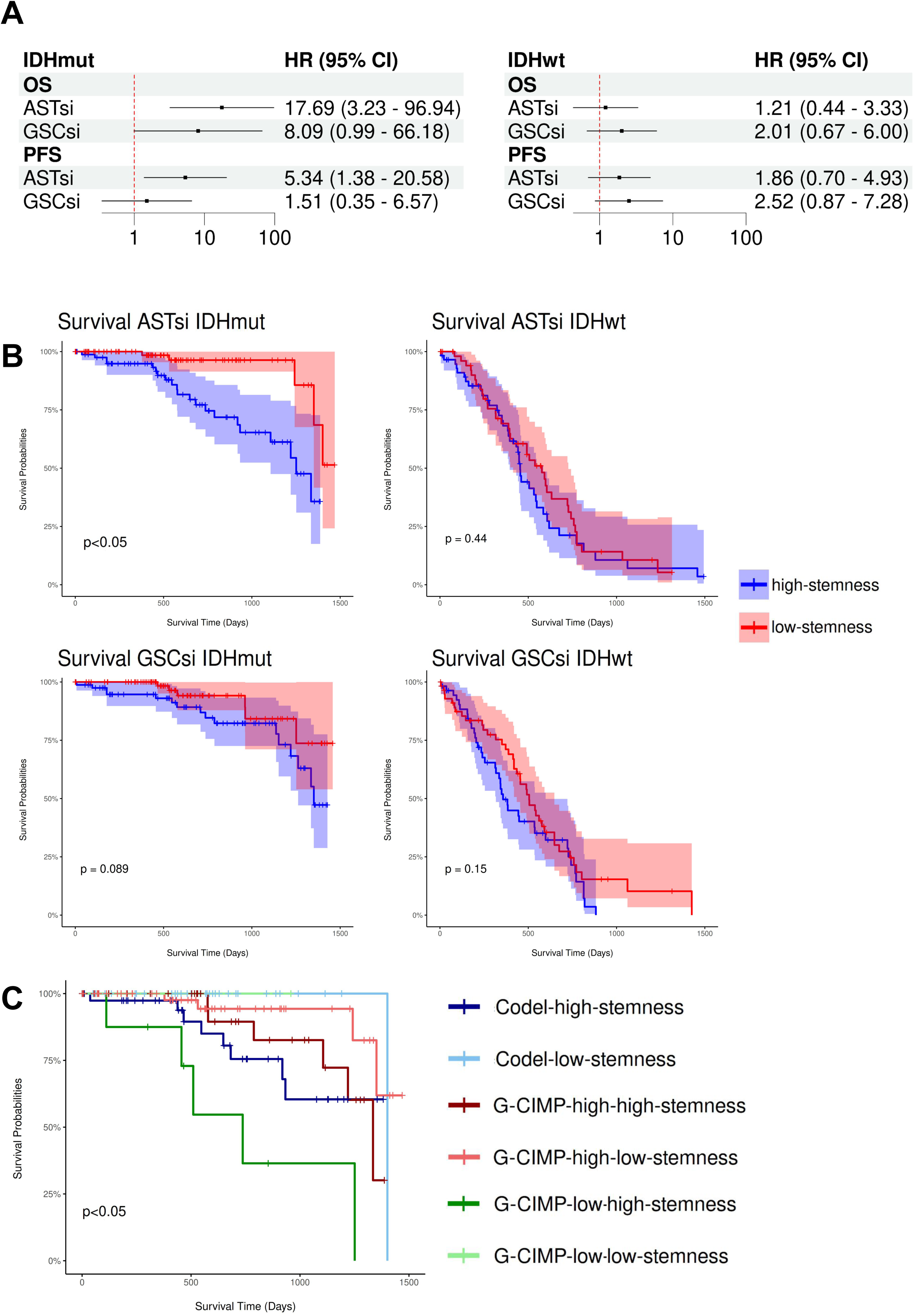
Survival Analysis Based on ASTsi and GSCsi in Glioma Patients from TCGA. A) Forest plot of Overall survival (OS) and Progression Free survival (PFS) of IDHmut (left, n=348 bulk RNA-seq samples) and IDHwt (right, n=229 bulk RNA-seq samples) glioma patients from TCGA associated with ASTsi and GSCsi. B) Kaplan-Meier curves comparing OS between High-stemness and Low-stemness patients in LGG/GBM data from TCGA (IDHmut n=348 bulk RNA-seq samples; IDHwt n=229 bulk RNA-seq samples). High-stemness defined as samples with ASTsi/GSCsi above the third quartile, that is, the 25% with the highest indices. Low-Stemness defined as samples with ASTsi/GSCsi below the first quartile, that is, the 25% with the lowest indices. In the left column the IDHmut samples (n=87 high-stemness, 87 low-stemness) and on the right the IDHwt samples (n=58 high-stemness, 58 low-stemness). C) Kaplan-Meier curves comparing OS between High-ASTsi and Low-ASTsi patients in TCGA IDHmut samples (n=348 bulk RNA-seq samples). Here the samples are divided by the methylation subtype defined by Cecarelli et. al, 2018. High-stemness defined as samples with ASTsi above the third quartile, that is, the 25% with the highest indices (n=42 high-ASTsi-Codel, 33 high-ASTsi-G-CIMP-high, 8 high-ASTsi-G-CIMP-low). Low-Stemness defined as samples with ASTsi below the first quartile for each methylation subtype, that is, the 25% with the lowest indices (n=32 low-ASTsi-Codel, 55 low-ASTsi-G-CIMP-high, 1 low-ASTsi-G-CIMP-low).

### 3. Genes related to Stemness and potential therapeutic targets

We then investigated the genes whose expression have the highest correlation with ASTsi or GSCsi to identify potential biomarkers. The ASTsi top 5 correlated genes (COPS3, OLA1, TIPRL, MRP35, and ORC5) showed hazard ratios greater than 1 in overall survival, and its high expression is directly related to lower survival in at least one of the IDHmut or IDHwt subtypes of gliomas (FIGURE 3A, FIGURE 3B). Notably, ORC5 showed evidence of high OS and PFS, and high expression of these genes indicates lower survival in both IDHmut and IDHwt gliomas. Regarding the GSCsi top 5 associated genes (SEPTIN6, SLC9A1, RASSF5, DYNLT3, and GPAT3), most of them presented a risk association considering OS and PFS. The only one to present evidence of direct association with worse survival separating the IDHwt and IDHmut samples was the SEPTIN6 gene, which showed worse survival in the IDHmut samples when it is more expressed (FIGURE 3A, FIGURE 3C).

**Figure 3:**
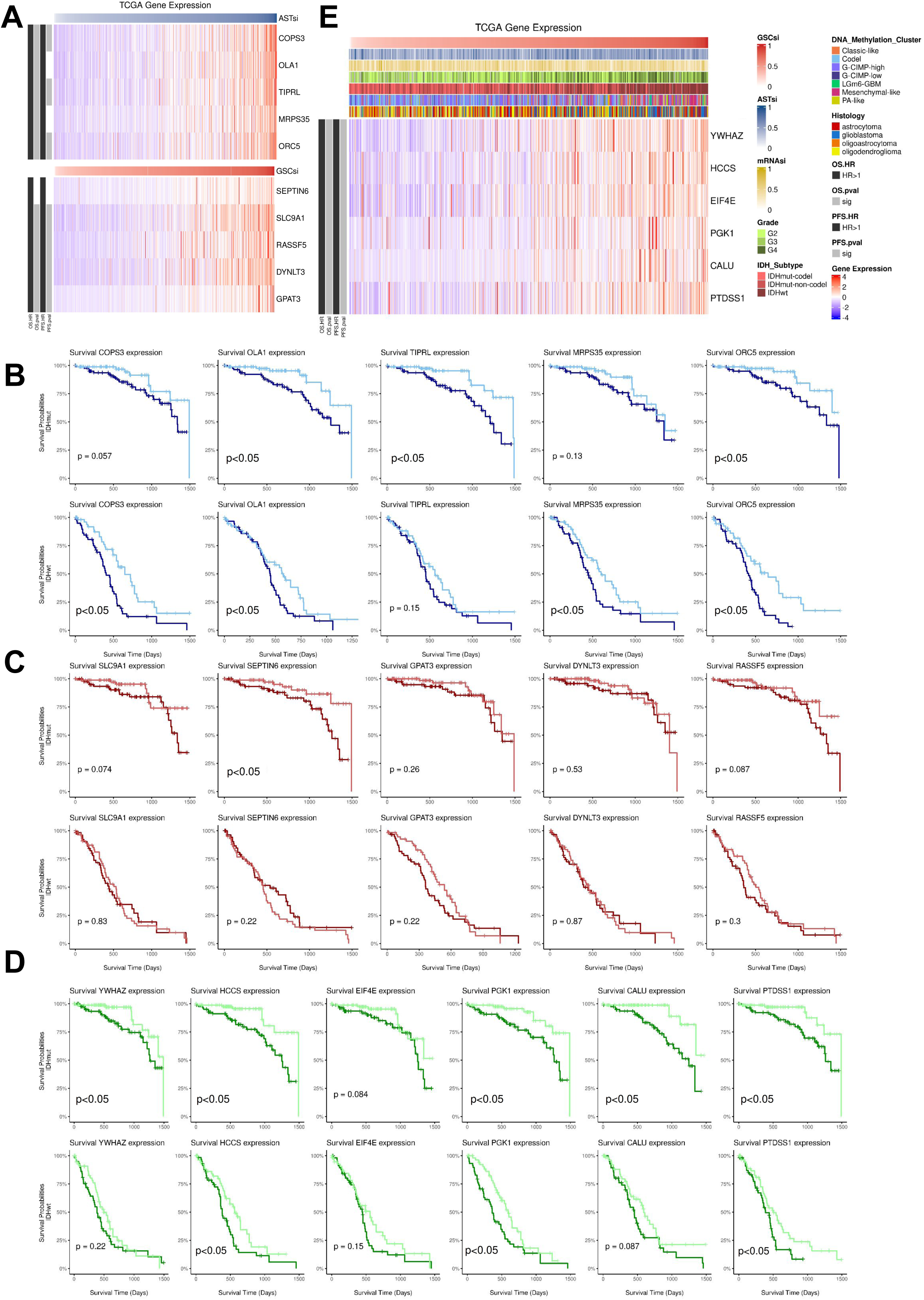
Gene Correlation and Survival Analysis in LGG/GBM Data from TCGA. A) Heatmap showing the genes that correlated with ASTsi and GSCsi simultaneously (p.value <0.05 and correlation>0.5) in the LGG/GBM data from TCGA (n=673 bulk

We also observed genes that simultaneously had a positive correlation greater than 0.5 with ASTsi and GSCsi. With these parameters, six common stemness related-genes were found between ASTsi and GSCsi: YWHAZ, HCCS, EIF4E, PGK1, CALU, and PTDSS1. All of these genes showed high hazard ratios for overall survival and progression-free survival within the TCGA cohort. High expression of HCCS, PGK1, and PTDSS1 was also associated with lower survival in both IDHmut and IDHwt samples (FIGURE 3D, FIGURE 3E).

These results highlight the importance of the discussed genes. For example, YWHAZ has been recognized as a potential biomarker in various types of cancer, playing a crucial role in diagnosis, prognosis, and even chemotherapy resistance ^11 12^. Similarly, HCCS expression has been associated with different clinical outcomes in various types of cancer, serving as a prognostic marker in renal cancer (with favorable effects) but unfavorable in liver and breast cancer, according to data available in the proteinatlas. PGK1, besides being related to chemotherapy resistance and prognosis in cancer patients in general, also shows potential as a therapeutic target in gliomas, where its regulation by miR-6869-5p influences cell proliferation and invasion ^13^. In contrast, elevated CALU expression is associated with more aggressive gliomas, suggesting a role in tumor progression, especially in higher grade and IDHwt glioma subtypes ^14^, and like PTDSS2, which has been identified as a possible key regulator in various types of cancer, including gliomas, perhaps PTDSS1, found in the results of this analysis, could also play an important role ^15^. These findings reinforce the functional diversity of these genes in different tumor contexts and highlight their potential as prognostic biomarkers and therapeutic targets in cancer, including gliomas.

### 4. Variation of ASTsi and GSCsi in longitudinal data

While TCGA is composed primarily of initial tumors prior to any treatment, the GLASS data mainly includes samples from patients who experienced recurrence. This approach allows us to compare results, check if the prediction models can identify characteristics such as IDH mutation and tumor grade in different cohorts, and analyze if the indices behave similarly in recurrent samples.

The prediction models were then applied to the gene expression data from GLASS. The indices obtained with the fetal astrocyte prediction model in primary tumors showed similar behavior to that found in TCGA data, considering grade and IDH status (Figure 4A). Gliomas grade IV and IDHwt showed decrease of ASTsi at recurrence, compared to the first surgery. Conversly, Gliomas IDHmut non-codels revealed an increase of ASTsi at recurrence, while no significant changes were observed for grades II and III or IDHmut codel. (Figure 4B). On the other hand, the GSCsi did not show evidence of differences considering the grade and IDH mutation of the primary glioma samples from GLASS (Figure 4C), which is different from what was observed in the TCGA data. While no GSCsi differences was observed between initial and recurrent samples for grades II and III and for IDHmut tumors, we found an increase in GSCsi when comparing samples from the first and second surgery in IDHwt and grade IV gliomas (Figure 4D).

**Figure 4:**
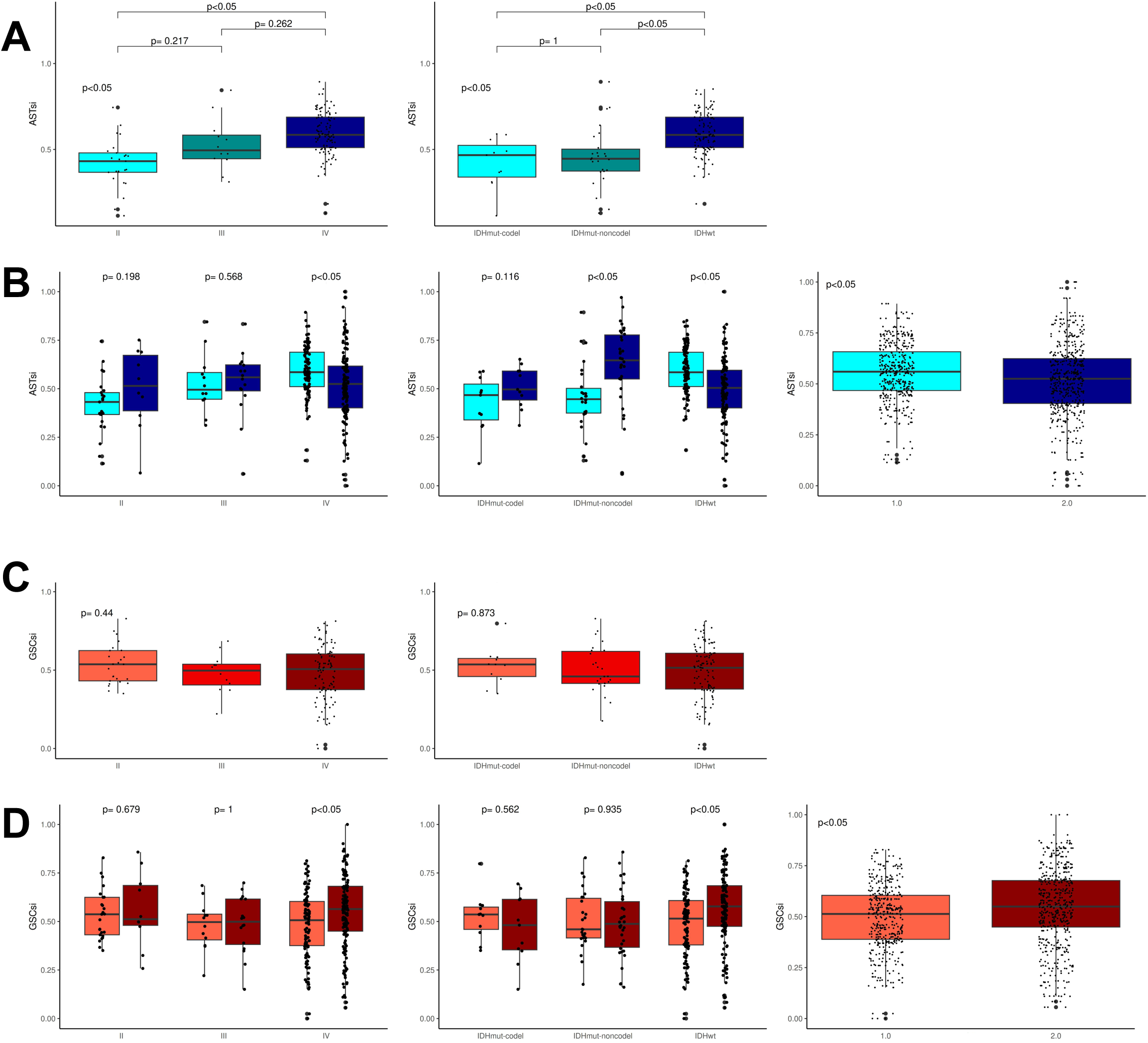
Analysis of ASTsi and GSCsi in Primary and Recurrent LGG/GBM Data from GLASS. A) Boxplots showing the distribution of ASTsi in primary LGG/GBM data from GLASS (n=144 bulk RNA-seq samples). On the left showing the distribution of ASTsi separating by tumor grade (n=25 G2, 12 G3, 106 G4), on the right showing the distribution of ASTsi separating by IDH mutation status (n=11 IDHmut-codel, 26 IDHmut-non-codel, 105 IDHwt). B) Boxplots showing the distribution of ASTsi in primaries and recurrents (second surgery) LGG/GBM data from GLASS (n=315 bulk RNA-seq samples), highlighting the difference in ASTsi between primary and recurrent tumors. On the left showing the distribution of ASTsi separating by tumor grade (n=25 primary and 10 recurrent G2, 12 primary and 16 recurrent G3, 105 primary and 144 recurrent G4), on the middle showing the distribution of ASTsi separating by IDH mutation status (n=11 primary and 11 recurrent IDHmut-codel, 26 primary and 34 recurrent IDHmut-non-codel, 105 primary and 124 recurrent IDHwt) and in the right showing the distribution of ASTsi between primary (n=144 bulk RNA-seq samples) and recurrent tumors (n=171 bulk RNA-seq samples) C) Boxplots showing the distribution of GSCsi in primary LGG/GBM data from GLASS (n=144 bulk RNA-seq samples). On the left showing the distribution of GSCsi separating by tumor grade (n=25 G2, 12 G3, 106 G4), on the right showing the distribution of GSCsi separating by IDH mutation status (n=11 IDHmut-codel, 26 IDHmut-non-codel, 105 IDHwt). D) Boxplots showing the distribution of GSCsi in primaries and recurrents (second surgery) LGG/GBM data from GLASS (n=315 bulk RNA-seq samples), highlighting the difference in GSCsi between primary and recurrent tumors. On the left showing the distribution of GSCsi separating by tumor grade (n=25 primary and 10 recurrent G2, 12 primary and 16 recurrent G3, 105 primary and 144 recurrent G4), on the middle showing the distribution of GSCsi separating by IDH mutation status (n=11 primary and 11 recurrent IDHmut-codel, 26 primary and 34 recurrent IDHmut-non-codel, 105 primary and 124 recurrent IDHwt) and in the right showing the distribution of GSCsi between primary (n=144 bulk RNA-seq samples) and recurrent tumors (n=171 bulk RNA-seq samples).

The results obtained when comparing indices in primary and recurrent samples provide evidence that IDHwt gliomas and grade IV tumors show a decrease in ASTsi and an increase in GSCsi upon recurrence. This change in stemness in more aggressive tumors may result from a phenotypic alteration of tumor cells during or after treatment, or an indication that gliomas with higher stemness related to GSCsi have greater resistance and recurrence capacity.

### 5. Analysis of GSCsi and ASTsi in scRNA-seq data of gliomas

Another aim of the prediction models is to serve as an alternative method for identifying potential GSCs in scRNA-seq data from glioma patients. The identification of GSCs or potential GSCs in gene expression data is still not a consensus, with multiple lists of marker genes or molecules that can be used in cytometry for cell isolation. To check if our prediction models are capable of identifying potential GSCs in scRNA-seq data, we used the core GBmap dataset, formed by the integration of 16 distinct scRNA-seq datasets of IDHwt gliomas (https://www.biorxiv.org/content/10.1101/2022.08.27.505439v1.full) and 3 distinct datasets of IDHmut gliomas (GSE70630, GSE89567, GSE102130).

In IDHwt samples, the distribution of GSCsi was more concentrated, while ASTsi appeared more dispersed and less specific (FIGURE 5A). Considering the annotation of tumor cells, it is noteworthy that GSCsi is higher in tumor cells, while ASTsi is higher in non-tumor cells (FIGURE 5B). Following the authors’ cell type annotation, we observed that malignant cells previously annotated as Stem-like and Differentiated-like have higher GSCsi compared to non-tumor cells, indicating that the GSC model is identifying tumor samples with higher indices and stem cell characteristics in scRNA-seq data (FIGURE 5C). On the other hand, ASTsi did not show a specific cell type for higher indices (not shown). Considering that these samples only encompass IDHwt gliomas, the result is expected since in previous analyses of this study, ASTsi was more effective in identifying aggressiveness and specific characteristics of IDHmut gliomas, while GSCsi was better associated with IDHwt gliomas.

**Figure 5:**
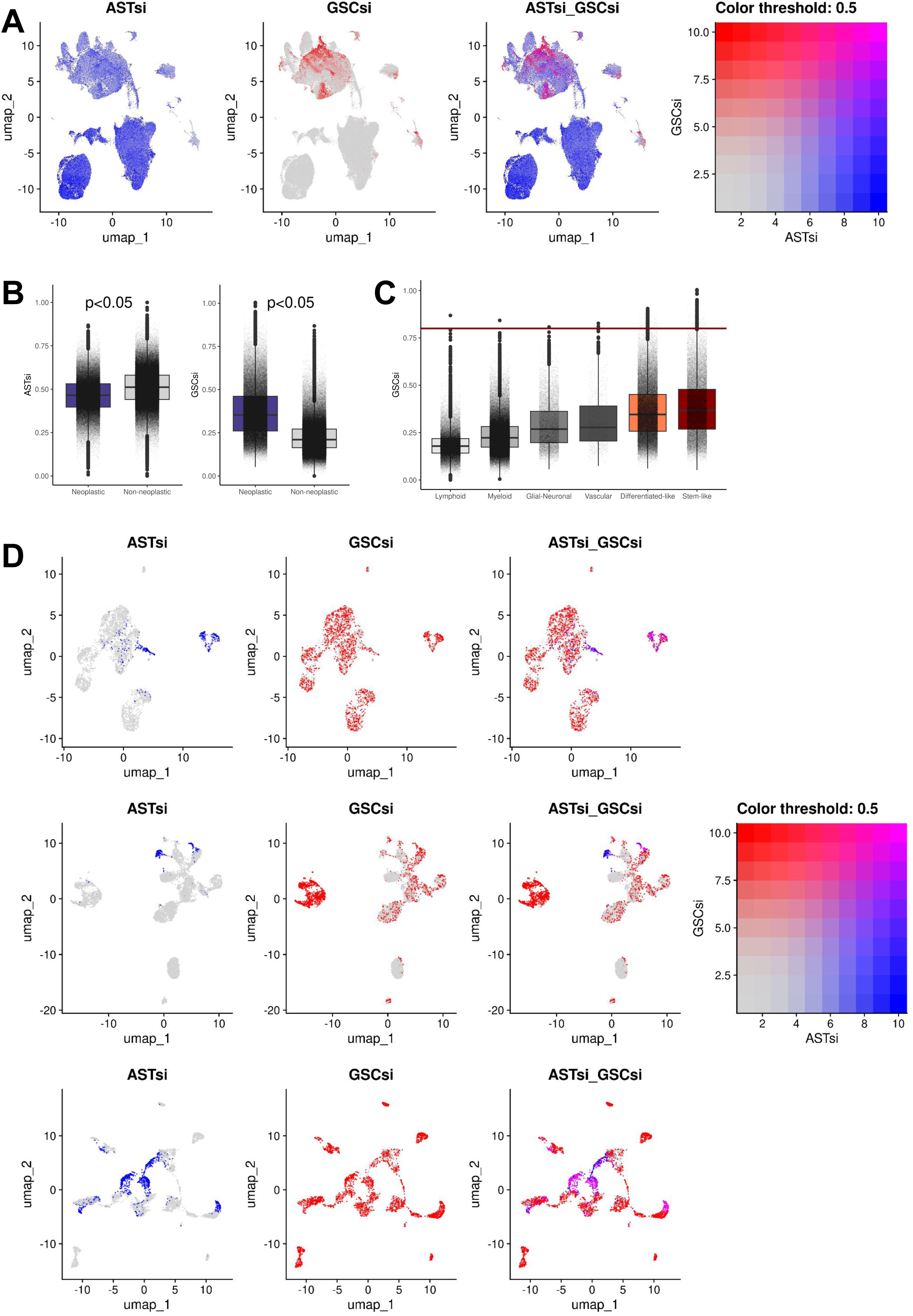
Stemness Indices analysis in Harmonized Glioblastoma and Glioma scRNA-seq Datasets. A) Uniform manifold approximation and projection (UMAP) plots of Human harmonized glioblastoma IDHwt scRNA-seq datasets (n=338564 scRNA-seq samples) highlighting in blue the samples with high ASTsi, in red the samples with high GSCsi and in pink the samples with high indices for both models simultaneously (continuous scale, from gray to the defined colors). B) Uniform manifold approximation and projection (UMAP) plot of Human harmonized glioblastoma IDHwt scRNA-seq datasets highlighting in purple on samples defined as tumor by the authors (GBmap methodology, referenced in table 1 of methods). On the right, boxplots showing the distribution of ASTsi and GSCsi in tumor (n=127521 scRNA-seq samples) and normal (n=211043 scRNA-seq samples) samples.

In IDHmut glioma datasets, it was observed that ASTsi was less dispersed than GSCsi regardless of tumor subclassification, which is opposite to what was found in IDHwt samples (FIGURE 5D). Interpretation regarding normal and tumor cells besides the definition of cell types was not performed in IDHmut datasets due to lack of information in the database and the raw expression matrix or FASTQ files not being provided by the authors. Although a few samples showed high indices for both models, the results indicate that ASTsi has a greater capacity to stratify and identify potential GSCs in IDHmut gliomas, while GSCsi shows better responses for IDHwt gliomas.

## Discussion / Conclusion

The results obtained reveal valuable insights into the biology of gliomas, highlighting the effectiveness of gene expression-based prediction models in identifying specific cellular subpopulations, such as fetal astrocytes and glioma stem cells (GSCs). A significant discovery is the association between histopathological and molecular characteristics of gliomas with the stemness indices generated by the prediction models. It was observed that the more aggressive subtypes of glioma present a higher presence of tumor stem cells, suggesting a relationship between tumor severity and the presence of these cells with greater potential for proliferation and resistance and these characteristics were captured by our prediction models.

The results of the survival analysis indicate that higher stemness, as indicated by ASTsi and GSCsi, is associated with a poorer prognosis for glioma patients, mainly in IDHmut samples. These findings suggest that the presence of tumor stem cells characterized by our models may be a relevant prognostic marker and a potential therapeutic target.

Investigation of genes correlated with ASTsi and GSCsi provided valuable insights into potential therapeutic targets. Genes such as YWHAZ, HCCS, PGK1, and CALU, which showed high correlation with stemness indices, are also associated with different clinical outcomes in various types of cancer, including gliomas, suggesting their potential as therapeutic targets.

Analysis of prediction models in longitudinal data showed variation in stemness indices between primary and recurrent tumors, especially in IDHwt gliomas and grade IV, highlighting the importance of monitoring and understanding these changes over time to develop more effective therapeutic strategies.

Finally, the application of the models in scRNA-seq data revealed their ability to identify potential GSCs in different subclasses of gliomas, offering a valuable tool for tumor heterogeneity analysis and identification of specific therapeutic targets.

In summary, gene expression-based prediction models represent a promising approach for characterizing and understanding the biology of gliomas, providing insights into their tumor heterogeneity, prognosis, and potential therapeutic targets. However, it is important to continue to refine these models and validate them on a large scale along with in vitro analyzes of patient cells or commercial cell lines for their effective clinical application.

## Materials and methods

**Table 1.** Resources Table.

| RESOURCE | SOURCE | IDENTIFIER |
| --- | --- | --- |
| Human iPSC-derived Astrocytes scRNA-seq data | Sloan, Steven A et al. 2017<sup>8</sup> | GEO database: GSE99951 |
| Progenitor and Mature Human Astrocytes bulk RNA-seq data | Zhang, Ye et al. 2015<sup>16</sup> | GEO database: GSE73721 |
| Human glioblastoma patient tumor scRNA-seq data | Odrzywolski A, Jarosz B, Kielbus M, et al. 2021<sup>9</sup> | European Genome-phenome Archive: EGAD00001006206 |
| Human iPS cell-derived neural stem cells bulk RNA-seq data | Tamura, Ryota et al. 2022<sup>17</sup> | GEO database: GSE150470 |
| Human glioma bulk RNA-seq data | TCGA database | https://portal.gdc.cancer.gov/ |
| Human glioma bulk RNA-seq data | GLASS database | https://glass-consortium.org/datasets/ |
| Human harmonized glioblastoma IDHwt datasets scRNA-seq data | Ruiz Moreno et al. 2022 (bioRxiv)<sup>18</sup> | https://cellxgene.cziscience.com/datasets |
| Human oligodendroglioma scRNA-seq data | Tirosh, Itay et al. 2016<sup>19</sup> | GEO database: GSE70630 |
| Human IDH-mutant astrocytoma scRNA-seq data | Venteicher, Andrew S et al. 2017<sup>20</sup> | GEO database: GSE89567 |
| Human K27M-mutant glioma scRNA-seq data | Filbin, Mariella G et al. 2018<sup>21</sup> | GEO database: GSE102130 |

## Data and Code Availability

This study did not generate new data. Table 1 describes the datasets used, including a brief description of each study, its source, and accession number. We accessed data available on NCBI using the ‘getGEO’ and ‘getGEOSuppFiles’ functions from the GEOquery package. The EGA dataset was accessed directly from the website after an access request. We accessed lower-grade glioma (LGG) and glioblastoma (GBM) data from TCGA using the ‘GDCquery’, ‘GDCdownload’, and ‘GDCprepare’ functions of the TCGAbiolinks package. After an access request, we accessed GLASS expression data. We accessed the Core GBmap scRNA-seq dataset directly via the CZ CELLxGENE link provided by the authors, as shown in the table. Details of the code and parameters used to create the prediction models including downstream analyses will be made available at https://github.com/RenanSimoesBR.

### Construction and Validation of Prediction Models

We built the prediction models using gene expression scRNA-seq data from fetal astrocytes derived from pluripotent stem cells (GSE99951) and derived gliomas stem cells (GSCs) (EGAD00001006206). We constructed individual Seurat objects for each dataset, excluding low-quality samples with fewer than 200 genes detected, genes not detected in at least 3 samples, and samples with more than 20% expression of mitochondrial genes. We normalized both Seurat objects using the standard Seurat pipeline with the ‘NormalizeData’, ‘FindVariableFeatures’, and ‘ScaleData’ functions.

To create the fetal astrocyte model (ASTsi), we selected samples from the cluster with the highest expression of fetal astrocyte marker genes (UBE2C, NUSAP1, TOP2A). For the glioma stem cell model (GSCsi), we selected glioma stem cells annotated by the original study, including GSCs derived from 3 glioblastoma patients (containing both whole tumor samples and GSCs). We used the normalized data from selected samples of fetal astrocytes and GSCs as input for the OCLR algorithm to create the prediction models, following the method described by Malta et al. (2018), using the ‘gelnet’ function of the gelnet package.

We validated the prediction models with external gene expression data from datasets containing samples of fetal astrocytes (GSE73721) and GSCs (GSE150470), along with their respective cell type annotations. We accessed the validation datasets in their already normalized format (TPM/FPKM) using the GEOquery package functions, and subsequently applied the prediction models.

### Survival Analyses

We performed survival analyses (Overall Survival, Progression-Free Survival, and Kaplan-Meier curves) on LGG and GBM data from TCGA, distinguishing between IDH wild type (IDHwt) and IDH mutant (IDHmut) samples. We obtained Hazard Ratios using the ‘coxph’ function of the survival package, applying age correction. We generated Kaplan-Meier curves for low-stemness and high-stemness groups using the ‘survfit’ function from the survival package and ‘ggsurvplot’ from the survminer package. Low- and high-stemness groups were divided based on quartiles. Samples with ASTsi below the first quartile were defined as low-ASTsi, and those above the third quartile were defined as high-ASTsi. The same parameters were used to define low-GSCsi and high-GSCsi.

### Statistical Analyses

For comparisons involving more than two groups, we used the Kruskal-Wallis method followed by Dunn’s post-hoc test with bonferroni correction to evaluate the statistical evidence of ASTsi and GSCsi among the different groups. For comparisons between two groups, we used the Wilcoxon test. We conducted correlation analyses between the obtained indices and/or gene expression using the Spearman method, which considers a monotonic relationship between variables, selecting samples with p.value<0.05. The integration of data from the 3 GBM patients with the derived GSCs was performed with the Seurat integration pipeline using the canonical correlation analysis (CCA) method. The specific processing, analysis and parameters used for scRNA-seq data with the Seurat package, as well as the R code, will be available on github https://github.com/RenanSimoesBR

**Table 2.** Packages Table.

| PACKAGE/RESOURCE | SOURCE | LINK/IDENTIFIER |
| --- | --- | --- |
| SeuratWrappers 0.3.5 | Community-provided curated by the Satija Lab | https://github.com/satijalab/seurat-wrappers |
| patchwork 1.2.0 | Thomas Lin Pedersen 2024 | https://github.com/thomasp85/patchwork |
| Seurat 5.1.0 | Hao, Y., Stuart, T., Kowalski, M.H. et al. 2024 | https://satijalab.org/seurat/ |
| glmGamPoi 1.6.0 | Constantin and Wolfgang 2020 | https://github.com/const-ae/glmGamPoi |
| GEOquery 2.62.2 | Davis, S. and Meltzer, P. S. 2007 | https://www.bioconductor.org/packages/release/bioc/html/GEOquery.html |
| stringr 1.5.1 | Hadley Wickham 2023 | https://CRAN.R-project.org/package=stringr |
| gelnet 1.2.1 | Artem Sokolov 2016 | https://CRAN.R-project.org/package=gelnet |
| ggpubr 0.6.0 | Alboukadel Kassambara 2023 | https://CRAN.R-project.org/package=ggpubr |
| clusterProfiler 4.2.2 | T. Wu 2021 | https://bioconductor.org/packages/release/bioc/html/clusterProfiler.html |
| TCGAbiolinks 2.22.4 | Colaprico A, Silva TC, Olsen C, et al. 2016 | https://bioconductor.org/packages/release/bioc/html/TCGAbiolinks.html |
| ggplot2 3.5.1 | H. Wickham 2016 | https://ggplot2.tidyverse.org/ |
| GEOquery 2.62.2 | Davis, S. and Meltzer, P. S. 2007 | https://www.bioconductor.org/packages/release/bioc/html/GEOquery.html |
| SummarizedExperiment 1.24.0 | Morgan M, Obenchain V, Hester J, Pagès H. 2024 | https://bioconductor.org/packages/release/bioc/html/SummarizedExperiment.html |
| reshape2 1.4.4 | Hadley Wickham 2007 | http://www.jstatsoft.org/v21/i12/ |
| survival 3.2.13 | Therneau T. 2021 | https://cran.r-project.org/web/packages/survival/index.html |
| ggfortify 0.4.17 | Yuan Tang, Masaaki Horikoshi, and Wenxuan Li 2016 | https://CRAN.R-project.org/package=ggfortify |
| circlize 0.4.16 | Gu, Z. 2014 | https://cran.r-project.org/web/packages/circlize/index.html |
| org.Hs.eg.db 3.14.0 | Marc Carlson 2021 | https://bioconductor.org/packages/release/data/annotation/html/org.Hs.eg.db.html |
| readxl 1.4.3 | Hadley Wickham and Jennifer Bryan 2023 | https://CRAN.R-project.org/package=readxl |
| dplyr 1.1.4 | Hadley Wickham, Romain François, Lionel Henry, Kirill Müller and Davis Vaughan 2023 | https://CRAN.R-project.org/package=dplyr |
| tidyverse 2.0.0 | Wickham et al. 2019 | https://www.tidyverse.org/ |
| rstatix 0.7.2 | Alboukadel Kassambara 2023 | https://CRAN.R-project.org/package=rstatix |
| survminer 0.4.9 | Alboukadel Kassambara, Marcin Kosinski and Przemyslaw Biecek 2021 | https://CRAN.R-project.org/package=survminer |
| gridExtra 2.3 | Baptiste Auguie 2017 | https://CRAN.R-project.org/package=gridExtra |
| ComplexHeatmap 2.10.0 | Gu, Z. 2016 | https://github.com/jokergoo/ComplexHeatmap |

## Notes

### Competing Interest Statement

The authors have declared no competing interest.

## Bibliography

1. Weller, M., Wick, W., Aldape, K., Brada, M., Berger, M., Pfister, S.M., Nishikawa, R., Rosenthal, M., Wen, P.Y., Stupp, R., et al. (2015). Glioma. Nat. Rev. Dis. Primers 1, 15017. 10.1038/nrdp.2015.17.

2. Wen, P.Y., Weller, M., Lee, E.Q., Alexander, B.M., Barnholtz-Sloan, J.S., Barthel, F.P., Batchelor, T.T., Bindra, R.S., Chang, S.M., Chiocca, E.A., et al. (2020). Glioblastoma in adults: a Society for Neuro-Oncology (SNO) and European Society of Neuro-Oncology (EANO) consensus review on current management and future directions. Neuro Oncol. 22, 1073–1113. 10.1093/neuonc/noaa106.

3. Hanahan, D. (2022). Hallmarks of cancer: New Dimensions. Cancer Discov. 12, 31–46. 10.1158/2159-8290.CD-21-1059.

4. Malta, T.M., Sokolov, A., Gentles, A.J., Burzykowski, T., Poisson, L., Weinstein, J.N., Kamińska, B., Huelsken, J., Omberg, L., Gevaert, O., et al. (2018). Machine Learning Identifies Stemness Features Associated with Oncogenic Dedifferentiation. Cell 173, 338–354.e15. 10.1016/j.cell.2018.03.034.

5. Lathia, J.D., Mack, S.C., Mulkearns-Hubert, E.E., Valentim, C.L.L., and Rich, J.N. (2015). Cancer stem cells in glioblastoma. Genes Dev. 29, 1203–1217. 10.1101/gad.261982.115.

6. Prager, B.C., Bhargava, S., Mahadev, V., Hubert, C.G., and Rich, J.N. (2020). Glioblastoma Stem Cells: Driving Resilience through Chaos. Trends Cancer 6, 223–235. 10.1016/j.trecan.2020.01.009.

7. Gimple, R.C., Bhargava, S., Dixit, D., and Rich, J.N. (2019). Glioblastoma stem cells: lessons from the tumor hierarchy in a lethal cancer. Genes Dev. 33, 591–609. 10.1101/gad.324301.119.

8. Sloan, S.A., Darmanis, S., Huber, N., Khan, T.A., Birey, F., Caneda, C., Reimer, R., Quake, S.R., Barres, B.A., and Paşca, S.P. (2017). Human Astrocyte Maturation Captured in 3D Cerebral Cortical Spheroids Derived from Pluripotent Stem Cells. Neuron 95, 779–790.e6. 10.1016/j.neuron.2017.07.035.

9. Couturier, C.P., Ayyadhury, S., Le, P.U., Nadaf, J., Monlong, J., Riva, G., Allache, R., Baig, S., Yan, X., Bourgey, M., et al. (2020). Single-cell RNA-seq reveals that glioblastoma recapitulates a normal neurodevelopmental hierarchy. Nat. Commun. 11, 3406. 10.1038/s41467-020-17186-5.

10. Ceccarelli, M., Barthel, F.P., Malta, T.M., Sabedot, T.S., Salama, S.R., Murray, B.A., Morozova, O., Newton, Y., Radenbaugh, A., Pagnotta, S.M., et al. (2016). Molecular profiling reveals biologically discrete subsets and pathways of progression in diffuse glioma. Cell 164, 550–563. 10.1016/j.cell.2015.12.028.

11. Yu, C.-C., Li, C.-F., Chen, I.-H., Lai, M.-T., Lin, Z.-J., Korla, P.K., Chai, C.-Y., Ko, G., Chen, C.-M., Hwang, T., et al. (2019). YWHAZ amplification/overexpression defines aggressive bladder cancer and contributes to chemo-/radio-resistance by suppressing caspase-mediated apoptosis. J. Pathol. 248, 476–487. 10.1002/path.5274.

12. Gan, Y., Ye, F., and He, X.-X. (2020). The role of YWHAZ in cancer: A maze of opportunities and challenges. J. Cancer 11, 2252–2264. 10.7150/jca.41316.

13. Zhang, K., Sun, L., and Kang, Y. (2023). Regulation of phosphoglycerate kinase 1 and its critical role in cancer. Cell Commun. Signal. 21, 240. 10.1186/s12964-023-01256-4.

14. Yang, Y., Wang, J., Xu, S., Shi, F., and Shan, A. (2021). Calumenin contributes to epithelial-mesenchymal transition and predicts poor survival in glioma. Transl. Neurosci. 12, 67–75. 10.1515/tnsci-2021-0004.

15. Yoshihama, Y., Namiki, H., Kato, T., Shimazaki, N., Takaishi, S., Kadoshima-Yamaoka, K., Yukinaga, H., Maeda, N., Shibutani, T., Fujimoto, K., et al. (2022). Potent and Selective PTDSS1 Inhibitors Induce Collateral Lethality in Cancers with PTDSS2 Deletion. Cancer Res. 82, 4031–4043. 10.1158/0008-5472.CAN-22-1006.

16. Zhang, Y., Sloan, S.A., Clarke, L.E., Caneda, C., Plaza, C.A., Blumenthal, P.D., Vogel, H., Steinberg, G.K., Edwards, M.S.B., Li, G., et al. (2016). Purification and Characterization of Progenitor and Mature Human Astrocytes Reveals Transcriptional and Functional Differences with Mouse. Neuron 89, 37–53. 10.1016/j.neuron.2015.11.013.

17. Tamura, R., Miyoshi, H., Imaizumi, K., Yo, M., Kase, Y., Sato, T., Sato, M., Morimoto, Y., Sampetrean, O., Kohyama, J., et al. (2023). Gene therapy using genome-edited iPS cells for targeting malignant glioma. Bioeng. Transl. Med. 8, e10406. 10.1002/btm2.10406.

18. Ruiz Moreno, C., Stunnenberg, H.G., Nilsson, M., Brandner, S., Kranendonk, M.E., Samuelsson, E., and Marco Salas, S. (2022). Harmonized single-cell landscape, intercellular crosstalk and tumor architecture of glioblastoma. BioRxiv. 10.1101/2022.08.27.505439.

19. Tirosh, I., Venteicher, A.S., Hebert, C., Escalante, L.E., Patel, A.P., Yizhak, K., Fisher, J.M., Rodman, C., Mount, C., Filbin, M.G., et al. (2016). Single-cell RNA-seq supports a developmental hierarchy in human oligodendroglioma. Nature 539, 309–313. 10.1038/nature20123.

20. Venteicher, A.S., Tirosh, I., Hebert, C., Yizhak, K., Neftel, C., Filbin, M.G., Hovestadt, V., Escalante, L.E., Shaw, M.L., Rodman, C., et al. (2017). Decoupling genetics, lineages, and microenvironment in IDH-mutant gliomas by single-cell RNA-seq. Science 355. 10.1126/science.aai8478.

21. Filbin, M.G., Tirosh, I., Hovestadt, V., Shaw, M.L., Escalante, L.E., Mathewson, N.D., Neftel, C., Frank, N., Pelton, K., Hebert, C.M., et al. (2018). Developmental and oncogenic programs in H3K27M gliomas dissected by single-cell RNA-seq. Science 360, 331–335. 10.1126/science.aao4750.

