## Supplemental Figure 1 for "Stemness prediction models reveal glioma aggressiveness and therapeutic targets in gliomas"

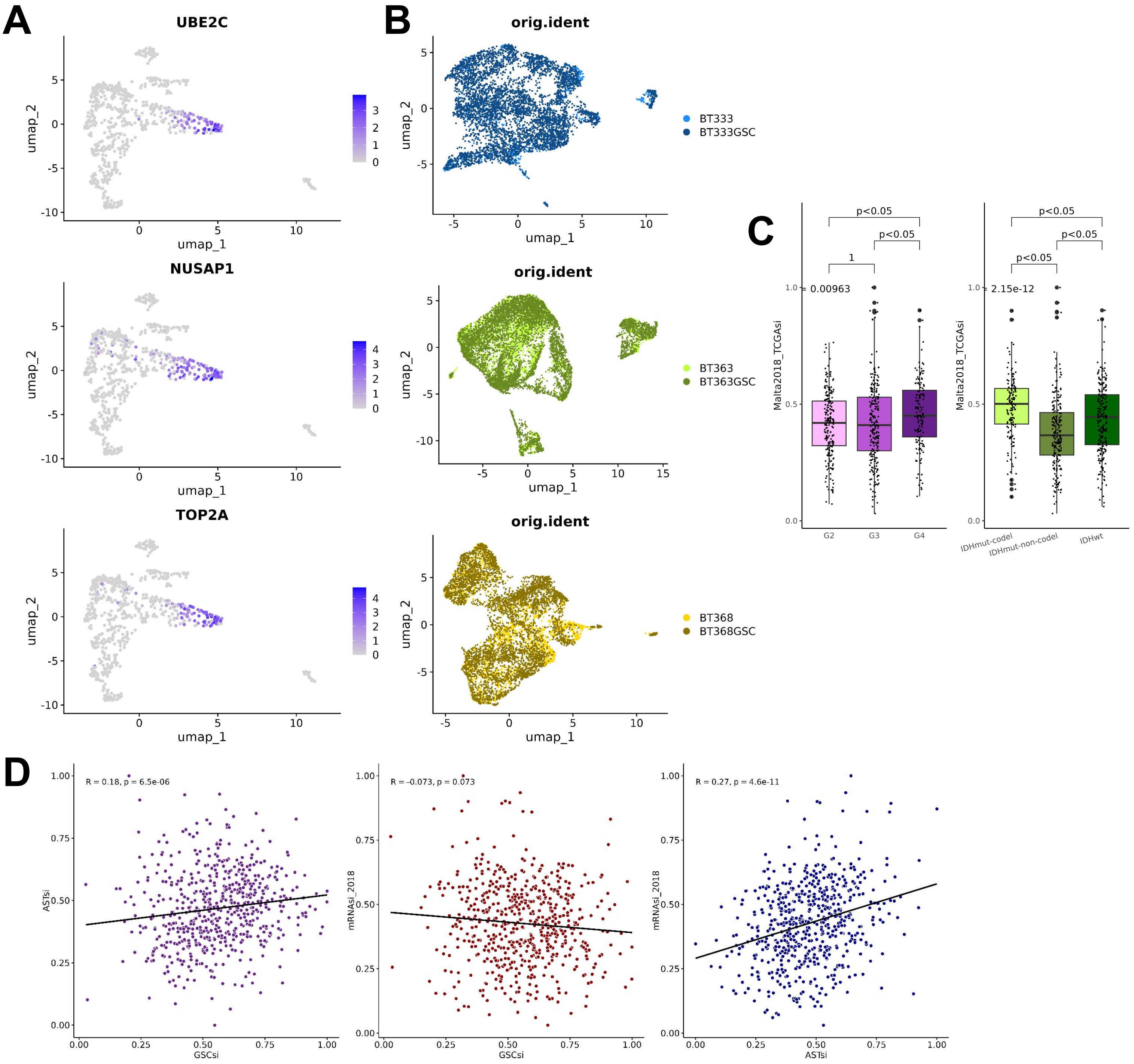
A
UBE2C
NUSAP1
TOP2A
umap_1
umap_2
B
orig.ident
BT333
BT333GSC
BT363
BT363GSC
BT368
BT368GSC
C
Malta2018_TCGAsi
0.00963
2.15e-12
p<0.05
1
G2
G3
G4
IDHmut-codel
IDHmut-non-codel
IDHwt
D
R = 0.18, p = 6.5e-06
R = -0.073, p = 0.073
R = 0.27, p = 4.6e-11
GSCsi
ASTsi
mRNAsi_2018

**Figure supp 1:**
A) Uniform manifold approximation and projection (UMAP) plots of Human iPSC-derived scRNA-seq samples (n=707 scRNA-seq samples) highlighting in blue the expression of marker genes for fetal astrocytes
B) UMAP plot of Human glioblastoma and derived GSCs scRNA-seq samples from the 3 patients that were used to construct the GSCs model after integration. In blue, samples from patient BT333 (n=614 whole tumor, 5146 derived GSCs scRNA-seq samples), in green, samples from patient BT363 (n=4334, 7872 derived GSCs scRNA-seq samples) and in yellow, samples from patient BT368 (n=2400 whole tumor, 4850 derived GSCs scRNA-seq samples)
C) Boxplots showing the distribution of mRNAsi_2018 in LGG/GBM data from TCGA (n=600 bulk RNA-seq samples). On the left showing the distribution of mRNAsi_2018 separating by tumor grade (n=214 G2, 241 G3, 145 G4), on the right showing the distribution of mRNAsi_2018 separating by IDH mutation status (n=151 IDHmut-codel, 226 IDHmut-non-codel, 223 IDHwt).
D) Scatterplot with Spearman correlation curve of the indices (ASTsi, GSCsi and mRNAsi_2018) obtained in the LGG/GBM samples from TCGA (n=600). In purple correlation between ASTsi and GSCsi, in red correlation between GSCsi and mRNAsi_2018, in blue correlation between ASTsi and mRNAsi_2018
